# Computational Screen for Sex-Specific Drug Effects in a Cardiac Fibroblast Network Model

**DOI:** 10.1101/2023.04.11.536523

**Authors:** Kelsey M. Watts, Wesley Nichols, William J. Richardson

**Affiliations:** Department of Bioengineering, Clemson University, Clemson, SC 29634, USA; Department of Chemical Engineering, University of Arkansas, Fayetteville, AR 72701, USA

## Abstract

Heart disease is the leading cause of death in both men and women. Cardiac fibrosis is the uncontrolled accumulation of extracellular matrix proteins which can exacerbate the progression of heart failure, and there are currently no drugs approved specifically to target matrix accumulation in the heart. Computational signaling network models (SNMs) can be used to facilitate discovery of novel drug targets. However, the vast majority of SNMs are not sex-specific and/or are developed and validated using data skewed towards male in vitro and in vivo samples. Biological sex is an important consideration in cardiovascular health and drug development. In this study, we integrate a previously constructed cardiac fibroblast SNM with estrogen signaling pathways to create sex-specific SNMs. The sex-specific SNMs maintained previously high validation when compared to *in vitro* experimental studies in the literature. A sex-specific perturbation analysis and drug screen uncovered several potential pathways that warrant further study in the pursuit of sex-specific treatment recommendations for cardiac fibrosis.

**Author Summary:** Heart failure is a leading cause of death for both men and women, but we still do not have adequate therapies to prevent or reverse this disease. One factor that contributes to heart failure is scarring of cardiac tissue, also known as fibrosis. Computer models can help find new heart failure drugs by simulating hundreds of biological reactions that regulate fibrosis at the molecular level. Unfortunately, the differences in male and female patients are not usually considered for these drug discovery simulations, which can result in drugs that work well for some individuals but not for other individuals. In our study, we added sex-specific biological reactions to a computer model in order to identify drugs that could treat fibrosis differently in male and female patients. Our simulations also predicted why premenopausal women may generally develop less fibrosis than men, while post-menopausal women may develop similar levels of fibrosis as men.

## INTRODUCTION

Heart failure (HF) currently afflicts roughly 6.2 million Americans and has a five-year survival rate of only 50% (1). Cardiac fibrosis, the uncontrolled accumulation of ECM proteins, can exacerbate the progression of HF (2–4). This deposition of ECM proteins is crucial to a patient’s initial survival of myocardial infarction (MI) to form the scar tissue that maintains the structural stability of the infarct region (5). However, when it continues uncontrolled, it becomes pathologic by reducing ejection fraction via thickening of the left ventricle wall and causing a greater chance of arrhythmia due to disruptions in electrical stimulus caused by collagen buildup (6,7).

The current treatment regimen for patients suffering from HF includes drugs such as angiotensin-converting enzyme (ACE) inhibitors and beta-blockers to reduce their blood pressure and slow their heart rate, which reduces stress on the heart in an attempt to slow the progression of HF (8). However, no current FDA-approved treatments directly target cardiac fibrosis (9). Ongoing research focuses on the development of a drug to directly inhibit cardiac fibrosis or even reverse it (10). Despite gains in understanding the complex signaling networks of cardiac fibroblasts - the drivers of cardiac fibrosis - finding new anti-fibrotic therapies has remained expensive and tedious, with low efficacy rates in clinical trials (11).

The use of computational models to conduct *in silico* simulations have been used in a variety of ways to accelerate drug discovery (12–15). Currently, these models are limited in scope and, at best, can narrow down a pool of potential drug candidates for further study. As the field of precision medicine grows, it is likely that the accuracy and predictive power of these models will also improve. However, in order to fully leverage the predictive power of these models, it is imperative that the data used to create them is robust. Unfortunately, experimental studies and clinical trials are historically skewed to male data. For several decades women of childbearing age were banned from participating in clinical trials (16). Even after this ban was reversed in the early 90s, it was not until 2016 that the National Institutes of Health required the use of both male and female animals in preclinical studies (17). This has resulted in several FDA-approved drugs having double the adverse drug response rate in women (18–20).

Biological sex is an essential factor when considering cardiovascular health (21–24). Most notably, premenopausal females are less likely to suffer from MI compared to age-matched men-primarily thought to be due to the cardioprotective role of the ovarian hormone estrogen (E2) (22). Additionally, the literature suggests that many adverse drug responses in both males and females occur in a sex-specific manner (25).

Including sex hormones into computational models could help correct for the dearth of sex-specific studies available in the literature and make precision medicine more accurate for patients. In this study, we aim to make a previously published signaling network model of cardiac fibroblasts sex-specific by incorporating E2 signaling. We then conducted a sex-specific drug screen to analyze divergences in drug response between males and females.

## MATERIALS AND METHODS

### Integration of estrogen into cardiac fibroblasts signaling network

A previously published signaling network model (SNM) of cardiac fibroblast was expanded to include the ovarian hormone estrogen (E2). The previous model was created via a manual literature search of ∼300 papers and included 125 nodes (i.e., proteins, integrins, cellular receptors, and transcription factors) and 174 edges (reactions) (26,27). Specifically, this model was comprised of 10 biochemical and biomechanical inputs, including transforming growth factor beta (TGFβ), angiotensin II (AngII), endothelin 1 (ET 1), and tension; downstream reactions of these inputs culminated in 22 cellular outputs, including alpha-smooth muscle actin (α-SMA), procollagen I & III (proCI and proCIII), several pro-matrix metalloproteinases and tissue inhibitors of metalloproteinases (proMMPs and TIMPs), and proliferation. New nodes and/or reactions were added to the SNM if two independent studies reported activation of inhibition of another node downstream of estrogen signaling. At least one of these two papers used for model advancement was from experiments conducted with cardiac fibroblasts; the second paper, if not from cardiac fibroblasts, typically reported results from experiments from other cardiac or fibroblast cell types. Nearly all of the papers used for model updates used neonate rat cardiac fibroblasts, pooling male and female cells together, as their cell type.

As previously described, the reactions of the SNM are governed by logic-based ODEs modeled as a system of Hill equations to capture node activity level (26–28). Logical NOT, AND, and OR gates were used for inhibitory and complex signaling interactions by applying logical operations: *f*_*INHIB*_*(x)=* 1-*f(x), f*_*and*_*(x*_*1*_, *x*_*2*_) = *f(x*_*1*_*)f(x*_*2*_), and *f*_*or*_*(x*_*1*_,*x*_*2*_*) = f(x*_*1*_*)+f(x*_*2*_*) - f(x*_*1*_*)f(x*_*2*_). Differential equations were constructed using the open-source software Netflux (https://github.com/saucermanlab/Netflux) for MATLAB (Mathworks, Natwick, MA) (28). Cytoscape was used to create all SNM visualizations included in this paper (29). All additional validation, perturbation, and drug screen simulations were conducted using MATLAB; finalized codes used in the analysis are available on (https://github.com/SysMechBioLab).

### Model Validation

Previous model validation of the cardiac fibroblast SNM conducted with 47 independent papers of direct measurement of model intermediate and output nodes found the SNM to be 81.8% accurate in predicting experimental activity levels of input-outputs (e.g., AngII treatment on proCI) and input-intermediates (e.g., AngII treatment out ERK activity level) found in the literature (26). In total, these 47 papers accounted for 118 perturbations of input outputs/intermediates validated by comparing literature experimental results to model predictions according to their change in activity level as ‘Upregulation’ (Δ Activity ≥5%), ‘Downregulation (Δ Activity ≤ -5%), or ‘No Change’ (−5% < Δ Activity < 5%).

To conduct a sex-specific validation of the updated model, 7 new papers were added to the validation set to account for estrogen and female-specific data. Each of the papers used for validation measured direct output secretion or intermediate signaling response (i.e., ELISA, IF, PCR, or Western Blot) to a single input stimulus in fibroblasts. In addition, previous perturbations were categorized based on the cell type used in analysis (male cells, pooled cells, or unreported; no previous papers used in the validation set reported data for female cells). Once disaggregated, the validation set contained 185 perturbation experiments which were 39% (n=73/185) male, 36% (n=66/185) pooled, and 17% (n=31/185) unreported, and 8% (n=15/185) female (Appendix S2).

Validation perturbations were grouped by experiments reporting results in female, male, and pooled neonate cells. Prediction accuracy was calculated using the same Δ Activity bounds described above using either the male or female SNM. To simulate the pooled neonate condition, model predictions from the male and female SNM were averaged prior to validation calculations. Model simulation predictions were generated in MATLAB by simulating basal conditions for 40 hrs, followed by simulating single input stimuli for 80 hrs. (26). An ideal EC_50_ of 0.65, n of 1.05, tension of 0.65, and input stimuli values of 1 were determined for each SNM through a parameter sweep (Appendix S3).

Additional validation was conducted to determine the accuracy of estrogen involvement in signaling pathways by comparing a combination of simultaneous treatments to experimental results. A tension weight of 0.65 was used and input stimuli weight of 1 for AngII, ET 1, and E2 was used to compare experimental results from Pedram et al. that reported E2 effect on AngII and ET 1 stimulation on α-SMA, fibronectin, proCI, and proCII in pooled neonate cardiac fibroblasts(30). A negative control simulation of input weights of 0.25 was generated by simulating basal conditions for 80 hrs, followed by simulating the treatment conditions (AngII, ET-1, AngII + E2, ET-1 +E2, and E2) for an additional 240 hrs.

### Network Perturbation Analysis

A network perturbation analysis was conducted in MATLAB as previously described to identify influential signaling nodes under different estrogen conditions (26). The input reaction weight of estrogen was attenuated, and all other input weights were kept at 0.4 for 80 hrs to simulate basal conditions. A Y_max_=0.1 knockdown of individual nodes for 240 hrs was used to identify sensitive nodes by calculating Δ Activity for each node as the sum of all knockdown simulation activity subtracted by basal conditions. Three perturbation conditions were conducted: 1) male (male SNM, E2 input w=0.25), 2) female post-menopausal (female SNM, E2 input w=0.5), and 3) female premenopausal (female SNM, E2 input w=1). Heat maps of the total knockdown effect on the SNM nodes were created and the top ten influential and sensitive nodes were determined for each condition.

### Drug Screen

In previous work, Zeigler et al. developed an approach to simulate the effects of known drugs on their cardiac fibroblast network model (12). In that prior study, 121 drugs were identified from DrugBank to connect with nodes in the model, totaling 36 unique drug-target interactions spanning agonists and antagonists, competitive and non-competitive (12,31). In our current study, we employed the same framework to perform a sex-specific drug screen using the three experimental conditions used for the perturbation analysis (male, female post-menopausal, and female pre-menopausal). A static application of drug administration (w=0.85) on profibrotic stimuli (all inputs except E2 w=0.4) for each of the three experimental conditions. To compare potential changes to cardiac fibroblast activity due to drug treatment, a Matrix Content Score was determined for each condition as MCS= Actvity_matrix_ + Activity_protease_ + Activity_inhib_ where Activity_matrix_ = (proCI +proCIII +fibronectin + periostin+osetopontin+LOXL1)/6; Activity_protease_ = (proMMPs 1, 2, 3, 8, 9, 12, 14)/7; and Activity_inhib_ = (TIMP1+TIMP2+PAI1)/3.

## RESULTS

### Integration of estrogen into cardiac fibroblasts signaling network

After a manual literature search of sex differences in cardiac fibroblasts, we used 18 previous reports to integrate E2 signaling into a published, large-scale signaling network model of cardiac fibroblasts via 7 new molecule nodes and 29 new reaction edges (Figure 1) (26,27,32– 49). In addition, 6 already present reactions were altered to include direct estrogen receptor inhibition activity. The added nodes include E2 and its three primary receptors: estrogen receptor alpha (ERα), estrogen receptor beta (ERβ), and g-protein coupled receptor (GPR30) (33,50). Cyclin beta 1 (Cyclinβ1) and cyclin-dependent kinase 1 (CDK1) were added downstream of GPR30 as they were not already included in the fibroblast SNM (39). The total network now consists of 132 nodes and 202 edges. The complete model, including all species (nodes), reactions, and default parameter settings, can be found in Appendix S1. In summary, default parameter settings included: reaction weights (w) as normalized activity levels between 0-1, y_int_=0, y_max_=1, and time constant (τ)= 1, 0.1, or 10 for signaling reactions, transcription reactions, and translation reactions respectively. Hill coefficient (n) and EC50 values for the new SNM were set at 1.05 and 0.65 respectively, based on a perturbation analysis. Sex-specific models were created by varying the Ymax of the three estrogen receptors (ER-β, ER-α, and GPR30) added to the model to mimic the physiological difference of varied expression and activity levels of estrogen receptors in male and female cells (33). The Ymax of the three estrogen receptors were set to 0.5 (50% of maximum saturation) and 1 (100% of maximum saturation) in the male and female SNM respectively.

**Figure 1:**
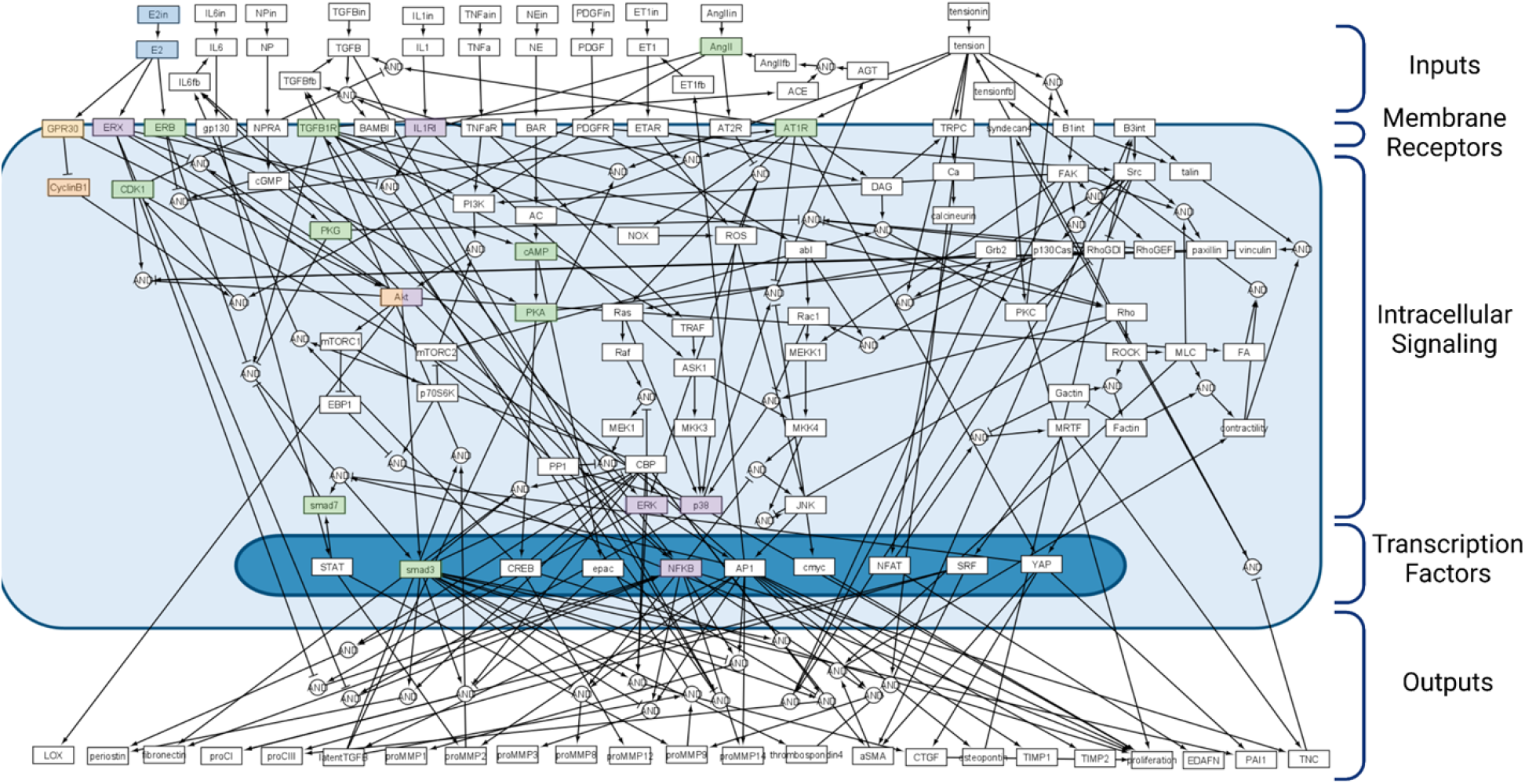
Cardiac fibroblast signaling network model (SNM) integrated with estrogen (E2) signaling. New nodes and their first downstream connections are highlighted in blue (E2), orange (GPR30), purple (ERα), and green (ERβ).

### Model accuracy remains high after integration of E2

To assess if model accuracy was influenced by sex, a sex-specific analysis was conducted using the sex-specific SNMs for model simulations compared with sex-specific experimental data reported in the literature (Figure 2). Previous perturbations used for model validation were categorized based on the cell type used in analysis (male cells, pooled cells, or unreported; no previous papers used in the previously published validation set reported data for female cells) (26). Additionally, 7 new papers were added to the validation set to account for estrogen and female-specific signaling (51–57). Once disaggregated, the validation set contained 185 perturbation experiments which were 39% (n=73/185) male, 36% (n=66/185) pooled, and 17% (n=31/185) unreported, and 8% (n=15/185) female. For the purpose of this paper, the 31 perturbations validated with experiments in the literature that did not report cell sex were not included in the rest of the analysis.

**Figure 2:**
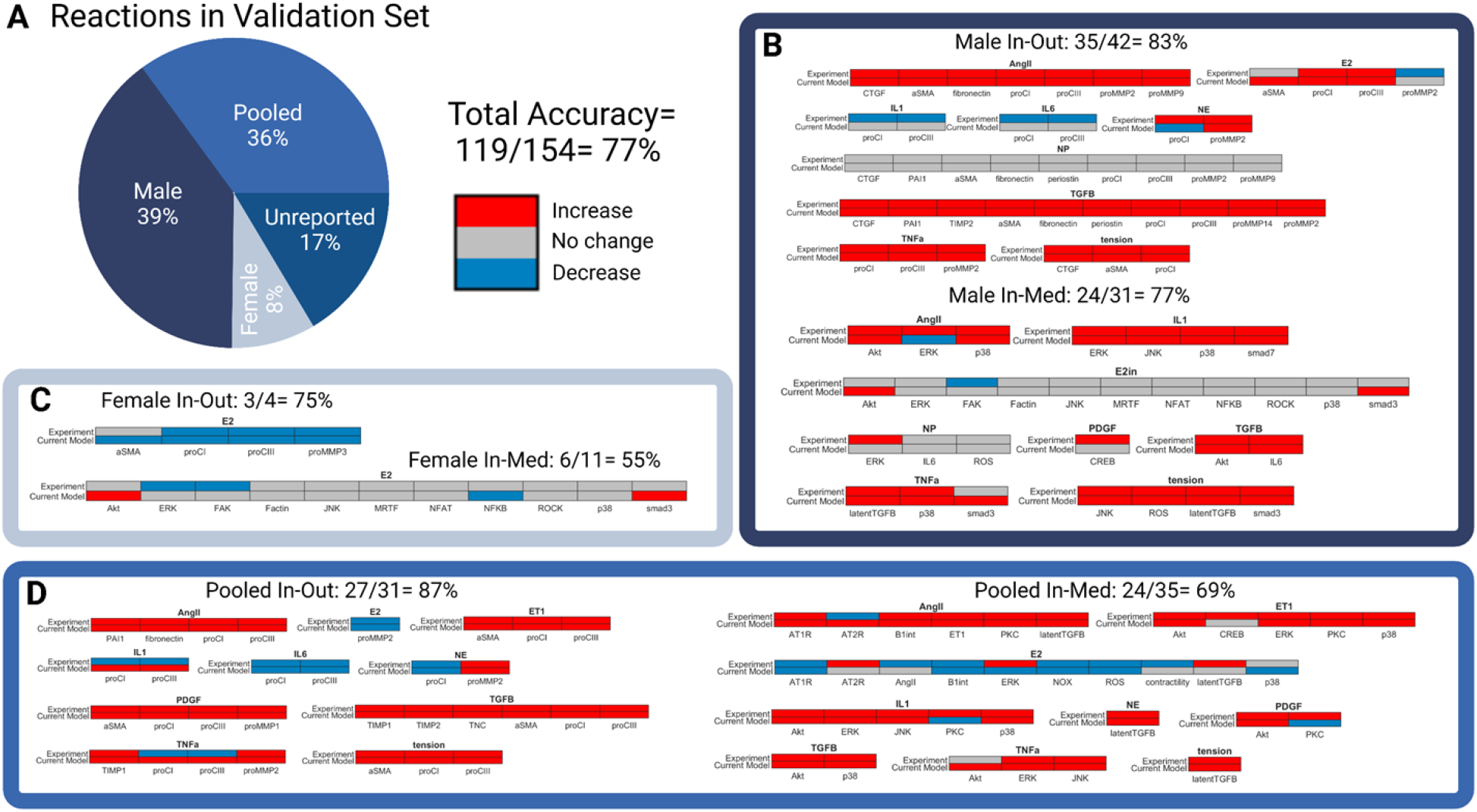
(A) Perturbation experiments used in previous signaling network model (SNM) validation were disaggregated by sex, and new reactions were added to the validation set to account for estrogen and female specific signaling. The reactions in the validation set comprised of 185 perturbation experiments which were 39% (n=73/185) male, 36% (n=66/185) pooled, 17% (n=31/185) unreported, and only 8% (n=15/185) female. For the purpose of this paper, the 31 perturbations validated with experiments in the literature that did not report cell sex were not included in the rest of the analysis. The total model accuracy of the new sex-specific SNM was determined to be 77%. (B-D) Individual biochemical stimuli sex-specific SNM predictions for input-output and input-intermediates compared to experimental studies found in literature in male (B), female (C), and pooled (D) cells. All model predictions were categorized as increase, no change, or decrease based on a ±5% change in activity levels compared to baseline conditions.

Accuracy of the model was relatively maintained after the integration of E2 in the SNM (compared to previous non sex-specific SNM accuracy of 81%) (26). When compared with 54 independent studies, 77% (119/154 simulations) of model predictions matched experimental results in the literature. Model predictions of estrogen treatment specifically were 59% (24/41 simulations) accurate across experimental conditions. Model predictions were the most accurate for the male SNM (81%, 59/73 simulations) and least accurate for the female SNM (60%, 9/15 simulations). The pooled SNM was 77% accurate (51/66 simulations). It is important to note that the female SNM model validation was only conducted with 15 simulations and compared with 3 papers. This is because very few papers report female-specific data for cardiac fibroblasts.

Notably, the model was accurate in predicting the divergent effect of estrogen treatment on male and female proCI and proCIII production, with E2 treatment increasing collagen production in males and decreasing it in females (51). Additionally, the inclusion of E2 into the model did not impact the accuracy of model predictions for other major cellular inputs, including AngII and TGFB. The lower accuracy of the model in predicting interleukin 1 (IL1), interleukin 6 (IL6), and neutrophil elastase (NE) effect on cellular outputs is consistent with the previously published SNM’s limitations (26).

### Model predicts E2 interactions with other input conditions

The experiments used in the validation set were all an analysis of a singular biochemical stimulus on cardiac fibroblasts. To determine if the model could accurately predict interactions between E2 and other fibrosis-driving input conditions, we compared simulation predictions against experimental data from Pedram et al. that investigated the effect of estrogen in conjunction with the fibrotic agonists angiotensin II (AngII) and endothelian-1 (ET-1) on pooled neonate cardiac fibroblasts (Figure 3) (30). The model predictions were consistent with *in vitro* results of the study, accurately predicting an increase in fibronectin, collagen 1, collagen III, and α-SMA production when stimulated with AngII or ET 1. Likewise, when these fibrotic agonists are paired with E2, the model predicts a return to control level values of the measured outputs. E2 stimulus alone also produced relatively little changes from control level predictions similar to experimental results. The only instance which the model did not qualitatively match experimental outputs, was that it failed to predict a substantial increase in α-SMA production for cells stimulated with ET-1 and therefore the effects due to combined treatment with ET-1 and E2 were minimal.

**Figure 3:**
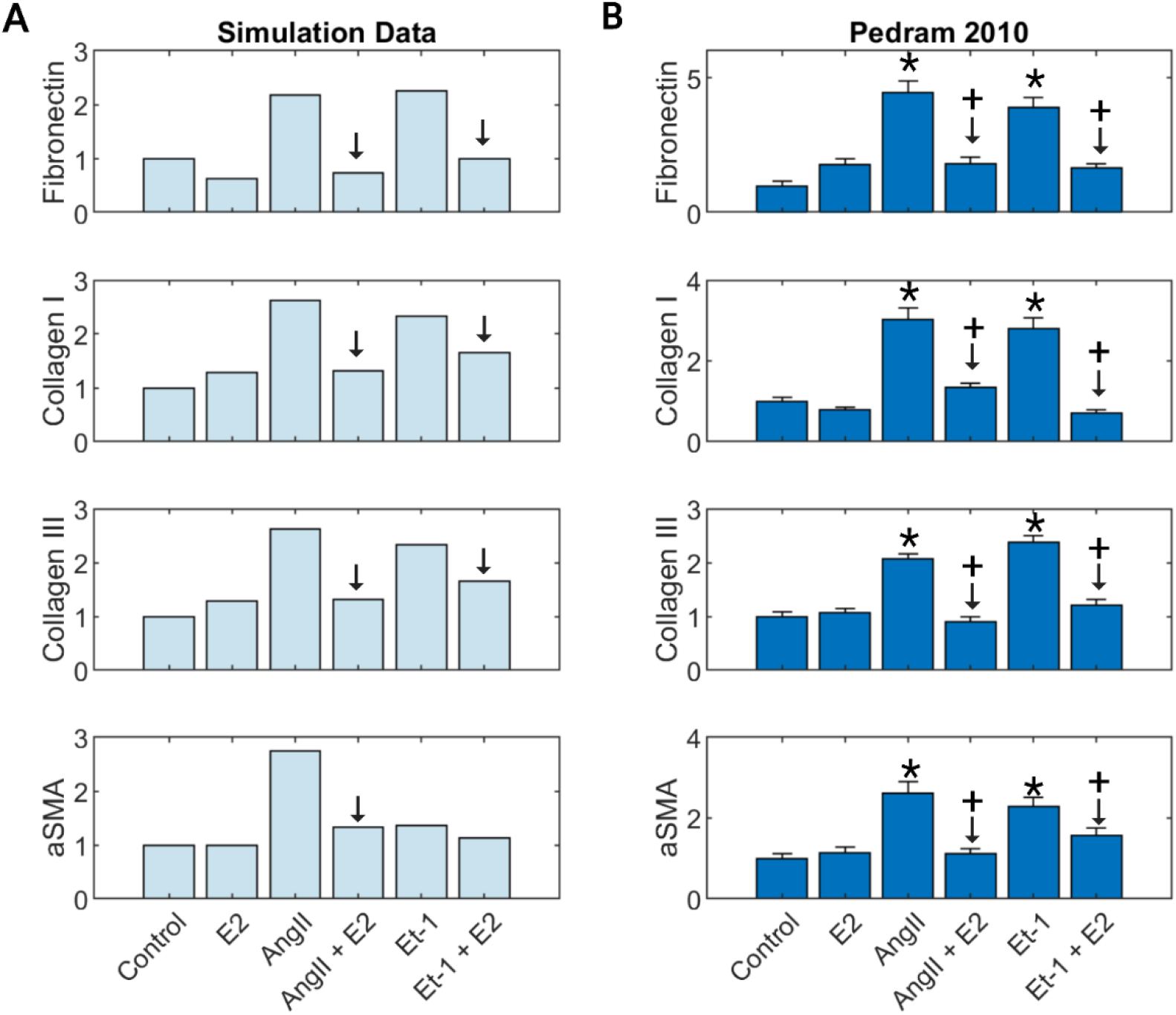
SNM predictions (A) compared to experimental *in vitro* results from the literature (B) for estrogen (E2) stimulus combined with fibrotic agonists angiotensin II (AngII) and endothelin 1 (ET-1) on fibronectin, collagen I, collagen II, and α-SMA production.

### Perturbation analysis identifies sex-specific network drivers and sensitivities

A perturbation analysis was conducted to computationally uncover the downstream effect of attenuating estrogen levels in the sex-specific SNMs (Figure 4). In attempts to mimic the physiological differences in estrogen receptor and circulating estrogen levels between females pre- and post-menopause and males, three input estrogen levels were used: 0.25 (25% saturation) for the male condition, 0.5 (50% saturation) for the female post-menopause condition, and 1 (100% saturation) for the female pre-menopause condition (23,58).

**Figure 4:**
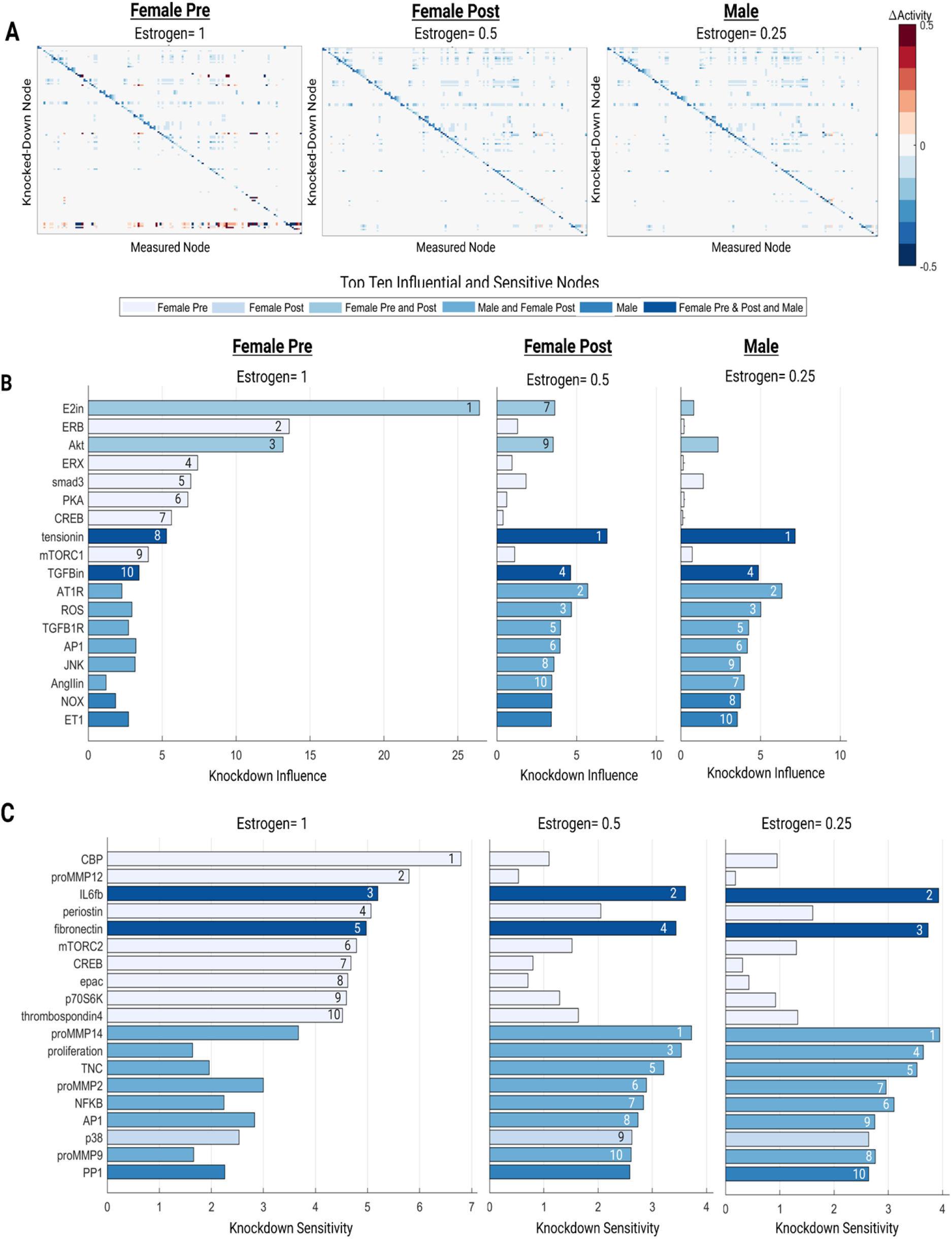
Total perturbation analysis for high (female pre-menopause), medium (female post-menopause), and low (male) estrogen conditions (A). Top ten influential (B) and sensitive nodes (C) for each experimental condition.

The top ten influential and sensitive nodes were determined for each condition (Figure 4B-C). Tension and TGFB were consistently in the top ten influential nodes for all conditions. ERB, ERX, smad3, PKA, CREB, mTORC1 were uniquely influential for the female pre-menopause condition, indicating potential pathways what may have a cardioprotective role downstream of estrogen. E2 and AKT were influential for both pre- and post-female menopause conditions. AT1R, ROS, TGFB1R, AP1, JNK, and AngII were influential in both the female post-menopause and male conditions. NOX and ET1 were only in the top ten influential nodes for the male condition indicating potential divergences in signaling pathways between male and female cardiac fibroblasts.

IL6 and fibronectin were sensitive across each estrogen perturbation. CBP, proMMP12, periostin, mTORC2, CREB, epac, p70S6K, and thrombospondin4 were in the top ten sensitive nodes only for the female pre-menopause condition. ProMMP14, proliferation, TNC, proMMP2, NFKB, AP1, and proMMP9 were sensitive in the female post menopause and male condition. Only p38 and PP1 were sensitive in just the female-post or male estrogen perturbations, respectively.

### Drug Simulation

A sex-specific drug screen (Figure 5) was conducted by perturbing nodes in the model corresponding to 36 unique drug targets or target combinations from DrugBank using a framework previously published by Zeigler et al (12). The effect of the drugs on fibrotic factors was analyzed when stimulated with various input stimuli with the same experimental conditions used in the perturbation analysis. The resulting changes in output expression levels indicate that drug effects tend to follow similar patterns for the post-menopausal female and male conditions, while the premenopausal female condition exhibits a more unique pattern. In particular, the premenopausal female model predicts more pro-fibrotic drug effects and fewer anti-fibrotic drug effects. This divergence is most dramatic in the case of NFKB + TNFa combinatory antagonism which predicts increases in matrix production and matrix content in the premenopausal female model but decreases in matrix production and matrix content in the post-menopausal female and male models. Individual treatments of TNFa antagonism or IL6 antagonism are also predicted to increase matrix content in premenopausal condition without affecting the other conditions. In total, six different drug strategies were predicted to be anti-fibrotic for post-menopausal female and male conditions with neutral or pro-fibrotic effects in the premenopausal case.

**Figure 5:**
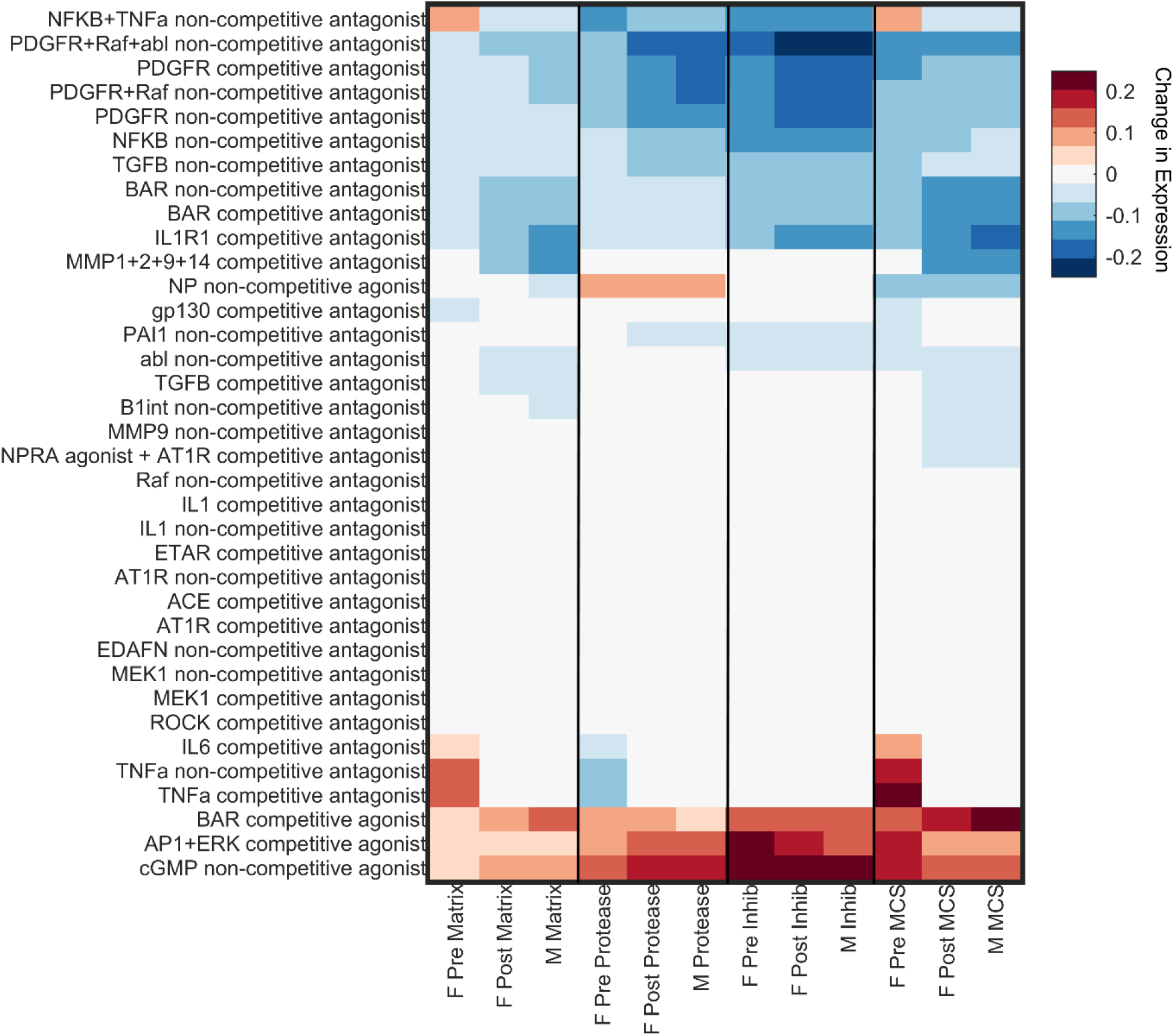
Drug screen analysis simulated the effects of 121 drugs with 36 unique drug-target interactions from DrugBank. Heatmap displays the average changes in protein output expression levels quantified as the difference between a baseline fibrotic condition and fibrotic condition + drug treatment. Outputs are indicated as ‘Matrix’ for matrix-related proteins (collagen I, collagen III, fibronectin, periostin, osteopontin, lysyl oxidase), ‘Protease’ for matrix-degrading proteins (MMP1, 2, 3, 8, 9, 12, 14), ‘Inhib’ for protease inhibitors (TIMP1, TIMP2, PAI1), and ‘MCS’ for an aggregate Matrix Content Score calculated as the average matrix - average protease + average inhibitor levels. Simulations were conducted for our three sex-specific modeling conditions: premenopausal female (‘F Pre’), post-menopausal female (‘F Post’), and male (‘M’).

## DISCUSSION

Developing a therapeutic strategy to target cardiac fibrosis has remained elusive due in part to the complex signaling mechanisms of cardiac fibroblasts, which can be influenced by biochemical factors, mechanical forces, and sex. In this study, we aimed to develop sex-specific network models of cardiac fibroblast signaling by integrating estrogen signaling into a previously published SNM of cardiac fibroblast biochemical and biomechanical pathways to understand sex-specific divergences in fibrotic signaling networks. Additionally, we conducted a sex-specific drug screen to demonstrate the model’s potential for facilitating sex-specific treatment recommendations.

Integration of estrogen into the SNM did not negatively alter its previously high predictive power in simulating experiments from the literature. The male SNM was the most accurate, which is unsurprising since the previously published versions of the model were built and validated using reported experiments that were overwhelmingly conducted with male cells (26,27). The accuracy of both the female SNM and the estrogen pathways were 15%-20% lower than the total SNM accuracy. This is likely because there are limited studies available in the literature that can be used to build and validate the model. Female data are consistently underreported in the literature, and even when female samples are included in analysis, almost no studies report the effect of fibrotic agonists on female cardiac fibroblasts outside the context of estrogen (59). Therefore, it limited our analysis of the female SNM accuracy in predicting the effect of the other model inputs such as AngII and TGFB on outputs and intermediates. The NIH requires use of both male and female cells and animals in funded research, however, the vast majority of studies that adhere to these guidelines do not disaggregate their results by sex (17,60). In the future, we recommend sex-specific reporting to help fill in the gaps of our understanding of sex-specific fibrotic signaling.

The results of the perturbation analysis uncovered potential regulatory nodes that are different between males and females pre- and post-menopause, which therefore warrant further study. Not only did these results highlight differences across distinct pathways, but several notable differences were even found across different isoforms in the same family. For example, the matrix metalloproteinase MMP12 exhibited heightened sensitivity in the premenopausal female condition, while MMP2, 9, and 14 sensitivities were heightened in the post-menopausal female and male conditions. MMPs are highly involved in fibrotic turnover with diverse functions involved in matrix proteolysis, cell adhesion, feedback signaling, and tissue mechanics, and there is emerging evidence of differences between male and female cells dependent on MMP type (61–64). Model predictions identified numerous other proteins that were sensitive to the different estrogen perturbations, but there is currently a lack of sufficient experimental data to determine if these findings are valid *in vitro* or *in vivo*.

The sex-specific drug screen clearly exhibited instances of varied drug response due to higher estrogen levels in the female pre-menopause SNM. For example, NFKB + TNFa combinatory antagonism predicted increases in matrix production and matrix content in the premenopausal female model, but the opposite response in the post-menopausal female and male models. The drug NFKB + TNFa combinatory antagonism was modeling was Thalidomide, a drug that in the 1950s caused severe birth defects in thousands of children after expecting mothers took it to alleviate morning sickness (65). More recently studies have exhibited the potential use of Thalidomide to prevent excessive cardiac remodeling in heart failure, but to our knowledge no studies have investigated the role E2 may play in negating its potential benefits (66,67).

The differences between the male and female post-menopause SNMs were much more subtle compared to the female-pre SNM. Although there were slight differences in the intensity of the effect of the drugs in up or downregulating fibrotic factors in a sex-specific manner, the results did not provide enough evidence to elucidate mechanisms that account for the adverse drug responses that are more likely to occur in females compared to men. A study that investigated the effect of age on adverse drug response in both men and women found that adverse events were the highest for women aged 30-59, but that adverse events being higher in women than men persisted throughout life (68). Although our drug screen does capture this likelihood of higher adverse effects in premenopausal women, it is possible that the integration of estrogen signaling alone was not enough to account for the sex-specific differences in fibrotic response to different therapeutics between the male and female postmenopausal models.

Many of the limitations of the sex-specific models designed in this study are due to the lack of sex-specific data, particularly female data, available to build and validate the model. An additional limitation of the current models is their lack of quantitative data to infer input parameter values. Without these inputs, the models can only make semiquantitative predictions that can be qualitatively compared to experimental literature but have little clinical significance. Because of this constraint, the model was only validated with estrogen levels of high (female pre-menopausal), medium (female post-menopausal), and low (male). However, like other hormones, estrogen levels can fluctuate from person to person and throughout life. Additionally females undergoing menopause are often considered perimenopausal for several years (median of 4) as hormone levels gradually change (69). These attenuations were not accounted for and would likely play a role in the downstream signaling response. Future studies could address this by testing with more levels of estrogen and integrating feedback loops for estrogen and its receptors.

A final limitation of the sex-specific SNMs was the use of only estrogen signaling to model sex differences. Not only are their additional sex hormones that could be incorporated in the future, there are also known biological sex dimorphism that are independent of gonadal hormones. Research shows that genetic differences between males and females can also contribute to cardiovascular disease presentation and severity (70,71). Future studies could address this by integrating genomic or transcriptomic data into the model to account for additional differences due to biological sex in addition to gonadal hormones (72).

## CONCLUSIONS

We integrated estrogen signaling into a cardiac fibroblast SNM in order to make sex-specific predictions of cardiac fibrosis. These models were validated to be 77% accurate in predicting experimental outcomes from the literature, making it a valuable tool to further study the sex dimorphisms in cardiac fibroblasts. Estrogen’s cardioprotective effects were evident in the drug screen of the premenopausal SNM. Differences between the male and female premenopausal conditions perturbation analysis and drug screen indicate several potential regulatory mechanisms that warrant further study. Future model development and validation will require more generation of sex-specific data to further enhance modeling capabilities for clinically relevant sex-specific predictions cardiac fibrosis and treatment.

## Supporting Information

**Appendix S1**. Fibroblast Sex-Specific SNM

**Appendix S2**. Sex-Specific SNM Validation

**Appendix S3**. Parameter Sweep

## Acknowledgments

The authors gratefully acknowledge funding from the National Institutes of Health (HL144927) and an American Heart Association predoctoral fellowship (827397).

